# Phthalate Exposure Induces Inflammatory Signaling and Suppresses Mitochondrial Function in Marine Mammal and Human Cells

**DOI:** 10.64898/2026.02.09.704935

**Authors:** Elizabeth R. Piotrowski, Emily K. Lam, Diana D. Moreno-Santillán, Kaitlin N. Allen, Daniel E. Crocker, Anders Goksøyr, José Pablo Vázquez-Medina

## Abstract

This study investigated the transcriptional and bioenergetic responses to monoethylhexyl phthalate (MEHP) in primary fibroblasts derived from northern elephant seals (*Mirounga angustirostris*), common dolphins (*Delphinus delphis*), and humans, using RNA-seq, extracellular flux assays, and high-resolution microscopy of the mitochondrial reticulum. MEHP exposure did not induce cytotoxicity but triggered species-specific changes in gene expression and mitochondrial metabolism and morphology. Human cells showed the greatest transcriptional response, upregulating genes involved in detoxification, antioxidant, and inflammation while downregulating lipid metabolism pathways. The highest dose also decreased mitochondrial respiration and increased mitochondrial fragmentation, triggering a metabolic shift toward glycolysis. Elephant seal cells showed delayed glycolytic shifts, maintaining mitochondrial respiration and upregulating antioxidant, immune, and metabolic pathway genes through the highest dose. Despite mitochondrial fragmentation, they upregulated mitochondrial fusion/fission and trafficking genes. Dolphin cells exhibited the fewest changes in gene expression, mostly in hormone signaling and mitotic pathways. They showed dose-dependent declines in both respiration and glycolytic rates, even at the lowest concentration, yet maintained mitochondrial structural integrity while upregulating stress- and hypoxia-induced genes. These distinct strategies highlight species-specific susceptibility to toxicant-induced stress, offering new insights into how marine mammals respond to plastic-derived contaminants and reinforcing the need for species-specific ecotoxicological risk assessments.

## 1. INTRODUCTION

Recent studies show rising levels of microplastics—and their chemical additives, including phthalates—in human tissues (Nihart et al., 2025). Phthalates are widely used plasticizers that enhance the flexibility of consumer and industrial products; however, they are not chemically bound to plastics, which allows them to leach into the environment (Hauser & Calafat, 2005; Teuten et al., 2009). Di(2-ethylhexyl) phthalate (DEHP), a commonly used phthalate in medical devices, cosmetics, inks, and cleaning products, is hydrolyzed in the liver into mono-ethylhexyl phthalate (MEHP), its primary bioactive metabolite (Albro et al., 1984; Wormuth et al., 2006). DEHP and MEHP can accumulate in cell membranes, prolonging internal exposure and enhancing toxicity (Routti et al., 2021). MEHP interferes with multiple molecular pathways, particularly via nuclear receptors such as PPARα and PPARγ, which regulate lipid metabolism, energy balance, and inflammation (Hurst & Waxman, 2003). Dysregulation of these receptors promotes endocrine disruption and metabolic disturbance (Manteiga & Lee, 2016). MEHP also disrupts steroid hormone biosynthesis by inhibiting key enzymes involved in steroidogenesis (Craig et al., 2011; Wang & Tremblay, 2012), and induces oxidative stress by increasing reactive oxygen species production and impairing mitochondrial function (Wang et al., 2012; Kim et al., 2023). These effects promote cellular damage, inflammation, and dysregulated immune responses (Manteiga & Lee, 2016; Kuo et al., 2023).

While the effects of MEHP have been extensively studied in terrestrial animals, knowledge of its impact on marine mammals remains limited. MEHP and other phthalates have been detected in marine mammals worldwide (Hart et al., 2018; Routti et al., 2021; Rian et al., 2020; Xie et al., 2022; Andvik et al., 2024; de Lima et al., 2024; Lemos et al., 2024). As apex predators, marine mammals are vulnerable to bioaccumulating lipophilic toxicants, including phthalates (Zantis et al., 2021; Kashiwabara et al., 2023; Merrill et al., 2023, Joseph et al., 2026). Notably, marine mammals also possess unique physiological adaptations that enable them to manage oxidative stress associated with diving-induced ischemia/reperfusion (Allen and Vázquez-Medina, 2019). While terrestrial mammals experience oxidative damage following ischemia/reperfusion events, marine mammals counteract these effects through elevated antioxidant defenses (Wilhelm Filho et al., 2002; Zenteno-Savín et al., 2002; Vázquez-Medina et al., 2007, 2012; Allen et al., 2024). Recent studies suggest that marine mammal adaptations to cope with diving-induced oxidant stress may provide an advantage in dealing with toxicant-induced oxidative stress, as shown by distinct species-specific responses in pinniped and human skeletal muscle cells exposed to cadmium (del Águila-Vargas et al., 2020) and DEHP (Brassea-Pérez et al., 2024). While both California sea lion and human cells exhibited increased oxidant generation when treated with DEHP, human cells experienced oxidative damage, cell death, and upregulated phase II detoxification processes, whereas sea lion cells activated antioxidant defenses and repair mechanisms (Brassea-Pérez et al., 2024). In contrast, recent evidence suggests that phthalates act as both agonists and antagonists for PPARG, glucocorticoid, and thyroid hormone receptors in fin whale fibroblasts (Mukundan et al., 2026, Preprint). While studies on phthalate toxicity in marine mammals are beginning to emerge, a comprehensive understanding of the cellular response to these compounds remains elusive. Here, we investigated changes in global gene expression, cellular energetics, and mitochondrial morphology in response to increasing concentrations of MEHP in primary dermal fibroblasts derived from common dolphins (*Delphinus delphis*), elephant seals (*Mirounga angustirostri*s), and humans.

## 2. MATERIALS AND METHODS

### 2.1. Animals

Northern elephant seal sampling was conducted under the National Marine Fisheries Service (NMFS) Permit No. 19108 (PI: Daniel Costa, UC Santa Cruz) at Año Nuevo State Reserve (San Mateo County, CA, United States). Primary cells were derived from seals and dolphins under NMFS Permit No. 22479. Seals were chemically immobilized via intramuscular injection of tiletamine-zolazepam, followed by intravenous administration of ketamine, as described previously (Vázquez-Medina et al., 2010). Skin biopsies (< 5mm) were obtained from the posterior flank region of each animal using a sterile scalpel and scissors. Dolphin tissue samples were obtained from necropsies of stranded long-beaked common dolphins at The Marine Mammal Center (TMMC; Sausalito, CA, United States). All tissue samples were rinsed with ice-cold sterile Hanks’ balanced salt solution (HBSS; Gibco, Thermo Fisher Scientific, Waltham, MA) containing 2% penicillin-streptomycin (P/S; Gibco), placed in fibroblast growth medium consisting of Dulbecco’s modified Eagle medium (DMEM; Gibco, catalog # 11885-084) supplemented with 10% fetal bovine serum (Seradigm, Avantor, Mexico), 10 mM HEPES (Gibco), and 1% antibiotic-antimycotic solution, and transported to the laboratory on ice.

### 2.2. Primary Cell Isolation and Culture

Dermal tissue samples were rinsed in ice-cold HBSS containing 2% penicillin-streptomycin (Gibco), minced into small pieces in a small volume of serum-free DMEM, and incubated (37°C, 5% CO2, humidified) in collagenase type II (0.66 mg/mL, Worthington, Danvers, MA) in DMEM (500 U/mL; Worthington Biochemical, Lakewood, NJ, United States) for 24 hours. Tissue suspensions were rinsed with HBSS, resuspended in fibroblast growth medium, and plated in tissue culture dishes coated with collagen II. Cell cultures were expanded and cryopreserved at passage 1. Primary dermal fibroblasts from three healthy adult human donors (passage 1) were obtained commercially (ScienCell Research Laboratories, catalog #2320) and cultured in commercial fibroblast medium (ScienCell, catalog #2301). Stock cultures from elephant seal (n = 3), dolphin (n = 2), and human (n = 3) cells were pooled together. To minimize the risk of phenotypic alterations associated with higher passage numbers and ensure passage consistency across species, we utilized cells at passages 4-7 for all experiments. Before conducting experiments, all cells were cultured in the same medium for 24 hours.

### 2.3. Immunofluorescence

Immunofluorescence was performed according to the protocol described by Vázquez-Medina et al. (2016). Briefly, cells were fixed with a 1:1 mixture of ice-cold acetone and methanol, followed by permeabilization, blocking, and overnight incubation with vimentin antibody diluted 1:100 (Cell Signaling Technology, catalog #5741). An Alexa Fluor 488 secondary antibody (Thermo Fisher) was used at a 1:200 dilution. Nuclei were counterstained with NucBlue for fixed cells (Invitrogen catalog #R37606). Controls lacking primary antibody were included in the experiments. Samples were visualized using a Zeiss Axio Observer 7 inverted microscope with a 20x objective and Zen software (Zeiss, Oberkochen, Germany).

### 2.4. MEHP Treatment

Mono(2-ethylhexyl)phthalate (MEHP, Sigma-Aldrich catalog #796832) was reconstituted in DMSO. Cells were treated with MEHP (0.5 *µ*M, 5 *µ*M, or 50 *µ*M) for 48h in complete medium. DMSO (0.1%) was used as a vehicle control. After treatment, cells were used for viability assays, RNA-seq, extracellular flux assays, or confocal microscopy analyses.

### 2.5. Cell Viability

Cell viability was measured using a RealTime-Glo™ MT Cell Viability Assay (Promega, catalog #G9711) and a SpectraMax M3 Multi-Mode Microplate Reader (Molecular Devices), according to the manufacturer’s protocol. Cells were treated with DMSO or MEHP for 48h as described above. Percent cell viability was calculated relative to untreated cells. Cells incubated with 1% Triton X100 were used as negative controls.

### 2.6. RNA-seq

Cells were lysed in RLT buffer without β-mercaptoethanol (Qiagen, Germantown, MD, USA). RNA was extracted using an RNeasy Mini Kit (Qiagen) with on-column DNase I digestion, following the manufacturer’s protocol, to remove genomic DNA contamination. RNA yield and purity were evaluated using a Nanodrop 1000 Spectrophotometer (Thermo Scientific). RNA integrity numbers (RIN) were determined using an Agilent 2100 Bioanalyzer (Agilent Technologies, Santa Clara, CA, USA). cDNA libraries were prepared from poly(A)-captured mRNA and sequenced on an Illumina® NovaSeq 6000 platform, yielding a total of 20 million paired-end 150 bp reads per sample, with estimated (Lander and Waterman, 1988) coverage depths of at least 2.8 times in elephant seal, 2.1 times in humans, and 2.4 times in common dolphin. Three replicates per condition (0.1% DMSO, 0.5 *µ*M MEHP, 5 *µ*M MEHP, 50 *µ*M MEHP).

### 2.7. Transcriptome Analyses

The quality of the raw reads was assessed with FastQCv0.12.1 (Andrews, 2010). Low-quality reads (PHRED score < 40) and adapter sequences were removed using Trimgalore v. 0.6.10 (Krueger, 2021). The remaining high-quality reads were mapped to the Northern elephant seal genome (GCA_021288785.2) annotated in (Torres-Velarde et al., 2021), the human genome (Ensembl: GRCh38 - hg38) or the common dolphin genome (NCBI RefSeq assembly: GCF_949987515.2) with STAR aligner v. 2.7.11a using default parameters (Dobin et al., 2013). Transcript quantification was performed using RSEM v.1.3.1 (Li and Dewey, 2011). Differential gene expression analysis was conducted using EBSeq with the rsem-run-ebseq wrapper, applying the multiple comparisons method (Leng et al., 2013) to evaluate treatments at 0.5 µM, 5 µM, and 50 µM MEHP relative to vehicle control. Differentially expressed genes were identified using a False Discovery Rate (FDR) threshold of 5%. Gene set enrichment analysis (GSEA) was conducted using normalized gene counts with GSEA software (Mootha et al., 2003; Subramanian et al., 2005). The analysis utilized the human hallmark gene sets (version 2024) from the Molecular Signatures Database (Subramanian et al., 2005; Liberzon et al., 2015). Gene sets were mapped using human gene symbols with the corresponding remapping chip (version 2024.1). The Signal2Noise metric was used for ranking, and statistical significance was determined through 1000 permutations. Pathways with an FDR below 25% were considered statistically significant.

### 2.8. Extracellular Flux Assays

Oxygen consumption rate (OCR) and extracellular acidification rate (ECAR) were measured in cells seeded in tissue culture-treated XF24 microplates using Seahorse Mitochondrial Stress Test Kits (Agilent Technologies, Santa Clara, CA, United States) and an XFe24 Extracellular Flux Analyzer. Optimal cell density and concentrations of oligomycin and carbonyl cyanide-p-trifluoromethoxyphenylhydrazone (FCCP) were determined empirically for each species; dolphin and seal cells were seeded at 40,000 cells/well, whereas human cells were seeded at 30,000 cells/well. Cells were washed with serum-free assay medium (Seahorse XF DMEM pH 7.4, 5.56 mM glucose, 1 mM pyruvate, 4 mM L-glutamine) and incubated at 37°C in a non-CO_2_ incubator for 1 h before the assay. OCR and ECAR were measured with oligomycin (2 *µ*M for human cells, 1.5 *µ*M for seal and dolphin cells), FCCP (2 *µ*M for human cells, 2 *µ*M for seal and dolphin cells), antimycin A (10 *µ*M), and 2-deoxyglucose (2DG, 50 mM for all cells). Total protein content was measured at the end of the experiment using the Pierce Rapid Gold BCA Protein Assay (Thermo Fisher Scientific). OCR and ECAR were normalized to total protein concentration.

### 2.9. Confocal Microscopy with 3D Image Reconstruction

Cells were grown in µ-Slide 8-well glass-bottom chambers (Ibidi), labeled with NucBlue, and transduced with CellLight Mito-RFP BacMam 2.0 fusion constructs (Life Technologies, cat #C10601) following the manufacturer’s guidelines and the signal was enhanced using a RFP-Booster Atto594 (ChromTek, cat, #rba59; 1:200) post-fixation. Cells were then fixed with 4% paraformaldehyde, rinsed in PBS, and imaged using a Zeiss LSM 880 FCS confocal microscope fitted with a 63X oil objective and Zen Black software (12 fields per condition for each species). *Z*-slices were collected and imported into Huygens Professional Software Scientific Volume Imaging B.V., The Netherlands, version 26.04) for deconvolution to reduce image distortion using the Classic Maximum Likelihood Estimation (CMLE) algorithm with the theoretical point-spread function, and a quality threshold of 0.01. Mitochondrial networks were analyzed using Imaris software (Oxford Instruments, Pleasanton, CA, version 10.2). Surfaces were added to the mitochondrial image to measure volume (µm^3^; the space a surface occupies) and disconnected components (the number of disconnected surfaces). At least 7 different cells per independent sample across all conditions and species were imaged and analyzed.

### 3.0. Western Blot

Cells were lysed in RIPA buffer containing 2% Halt Protease and Phosphatase Inhibitor Cocktail. Total protein concentration was determined using the BCA Rapid Gold Protein Assay Kit (Pierce Biotechnology). Proteins were diluted with lithium dodecyl sulfate buffer containing β-mercaptoethanol and denatured at 70°C for 10 min. 30 *µ*g of protein were resolved on 4-12% BT gels prior to transfer to nitrocellulose membranes. Membranes were incubated overnight with antibodies against mitofusin 2 (Mnf2; Cell Signaling Technology, cat #9482; RRID:AB_2716838; 1:1000). Proteins were visualized using IRDye 800CW secondary antibodies (LICOR) and a two-color near infrared system (Azure 500, Azure Biosystems). Membranes were stripped and reprobed with an antibody against α-tubulin (Cell Signaling Technology, cat #2144; RRID:AB_2210548_; 1:1000). Protein band intensity was quantified using FIJI v2.16.0 and normalized to α-tubulin.

### 3.1. Statistical Analyses

Statistical analyses and data visualization were conducted using R v4.1.2 in RStudio v2026.01.0+392 (Posit team, 2026). Normal distribution was assessed using Shapiro-Wilk tests, and equality of variances was assessed using Levene’s test via the *car* package (Fox et al., 2024). For within-species comparisons of 3D reconstructed mitochondrial network metrics (total network volume and the number of disconnected components), relative Mnf2/α-tubulin (change from control) levels, and extracellular flux assays, groups meeting all parametric assumptions were evaluated using a one-way analysis of variance (ANOVA) followed by Tukey’s Honestly Significant Difference (HSD) post-hoc testing using the R *stats* package (R Core Team, 2021). For datasets that violated these assumptions, non-parametric analyses were conducted using the Kruskal-Wallis test, followed by pairwise comparisons with the Wilcoxon ranking-sum test, with a Bonferroni p-value adjustment. For cell viability assays, data met all parametric assumptions and were evaluated using a one-way ANOVA, followed by pairwise t-test with Benjamini-Hochberg adjustment for multiple comparisons. Statistical significance was considered at *P* ≤ 0.05. Results are expressed as mean ± SE.

## 3. RESULTS AND DISCUSSION

### 3.1. Marine Mammal and Human Fibroblasts Proliferate in Primary Culture and Remain Viable after Exposure to MEHP

We established primary fibroblast cultures from dermal tissues of common dolphins and northern elephant seals and obtained primary dermal fibroblasts from humans to compare the effects of MEHP between marine mammal and human cells. Cells from all three species stained positively for the mesenchymal marker vimentin, confirming their fibroblast identity (Fig. 1A). We then tested whether 48 hours of MEHP treatment decreased cell viability. Cells from all species remained viable 48 h after MEHP treatment (Human: *P =* 0.71, Dolphin: *P =* 0.31, Seal: *P =* 0.62) (**Fig. 1B**), indicating that long-term MEHP treatment at our experimental doses exerts non-lethal effects in primary elephant seal, dolphin, and human dermal fibroblasts, consistent with previous studies (Giovani et al., 2022). Notably, exposure to the highest dose upregulated genes involved in cell cycle in cells from all three species (see Section 3.2), suggesting a coordinated stress response and possibly reflecting compensatory mechanisms that promote cell proliferation to avoid damage, preserve tissue function, or support repair, aligning with previous studies showing MEHP’s mitogenic effects and activation of proliferative signaling cascades (Luo et al., 2018; Zhou et al., 2023; Li et al., 2024). Notably, only 5 DEGs were shared across species at the high dose, suggesting species-specific responses (see Section 3.2).

**Figure 1.**
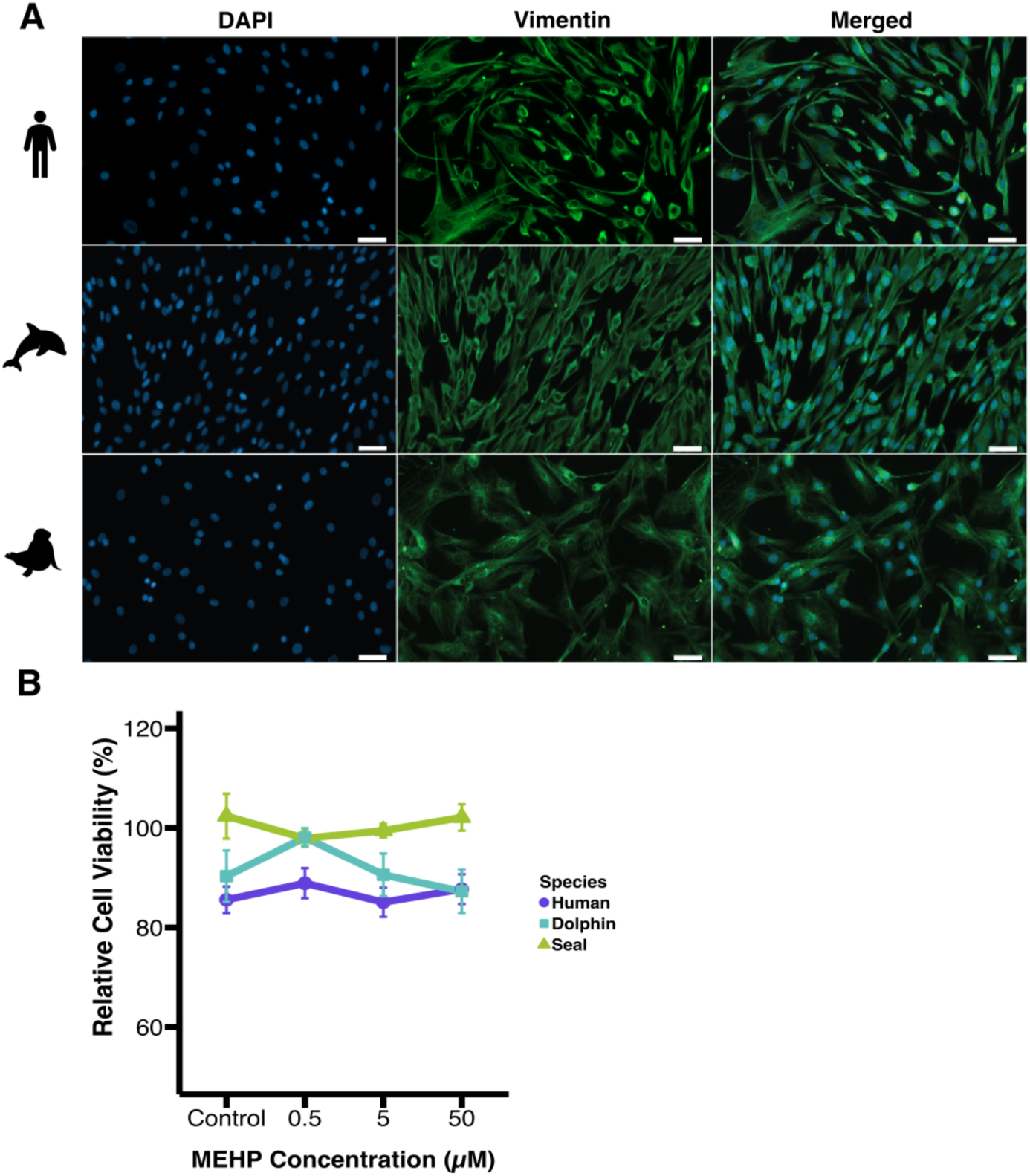
Primary fibroblasts derived from human, elephant seal, and dolphin skin stain positive for vimentin and remain viable after treatment with MEHP. (A) Immunofluorescence analysis for vimentin. Scale bar = 50 *µ*m. (B) Cell viability after treatment with increasing concentrations of MEHP for 48h. Results are expressed as means ± SE. N = 3-5 replicates per species per condition.

### 3.2. MEHP Exposure Differentially Regulates Gene Expression in Human, Elephant Seal, and Dolphin Cells

We used RNA-seq to evaluate global changes in gene expression in human, dolphin, and seal cells following 48 h incubation with three ascending concentrations of MEHP. We detected differentially expressed genes (DEGs) in response to MEHP treatment in cells from all species (**Figs. 2A-C**). Overall, human cells exhibited the largest shift in gene expression with a total of 2,183 (976 upregulated & 1,207 downregulated) DEGs (**Fig. 2A**). In contrast, dolphin cells were the least responsive with a total of 188 (60 upregulated & 128 downregulated) DEGs (**Fig. 2B**). Elephant seal cells showed more changes in gene expression compared to dolphin cells, but fewer changes compared to human cells, with 1,277 (643 upregulated & 634 downregulated) DEGs (**Fig. 2C**). These data align with previous work suggesting that phthalates can exert differential effects on mammalian cells, depending on the species, cell type, and level of exposure (Kluwe 1982; Ito and Nakajima, 2008; Rusyn et al., 2008).

MEHP upregulated and downregulated genes in a dose-dependent manner in all species, with the largest number of DEGs consistently observed at the high dose (50 *µ*M) (**Figs. 2A-2C**). At this dose, human cells showed the most DEGs, 749 upregulated and 975 downregulated (**Fig. 2A**), which is consistent with findings from previous studies reporting dose-dependent transcriptional shifts in human cells exposed to phthalates (Nazzari et al., 2023; Yasuda et al., 2024). In contrast, dolphin cells were the least responsive, with only 57 upregulated and 103 downregulated DEGs (**Fig. 2B**), suggesting lower sensitivity. At the low dose (0.5 *µ*M), elephant seal cells were the most responsive, with 190 upregulated and 204 downregulated DEGs (**Fig. 2C**), suggesting species-specific regulatory mechanisms among marine mammals. In contrast, dolphin cells had only 2 upregulated and 11 downregulated DEGs (**Fig. 2B**), and human cells had 21 upregulated and 30 downregulated DEGs at the low dose (**Fig. 2A**). At the medium dose (5 *µ*M), both human and elephant seal cells showed similar responses with 206 upregulated and 202 downregulated DEGs (**Figs. 2A, 2C**). In comparison, dolphin cells had only one upregulated and 40 downregulated DEGs (**Fig. 2B**).

Previous work from Fossi et al. (2025) linked phthalate exposure, including MEHP, to genes encoding key receptors (PPARA, THRA, THRB) and regulatory or structural functions (PKP3, PCBP3, SETD1A, OTX1, GRAMD4, CDC93) in fin whale skin. Though not statistically significant (FDR > 0.05), we observed similar trends in these genes in human, elephant seal, and dolphin fibroblasts. Together, these patterns may reflect potential conservation of these molecular targets across species, although the magnitude of their response may vary.

We next identified genes that were differentially expressed across all MEHP treatments in each species. We found 9 genes in human cells, 1 gene in dolphin cells, and 62 DEGs in seal cells, irrespective of the MEHP dose (**Fig. 2D-2F**). Of note, we did not find any DEGs shared amongst the three species in response to the low dose, 1 in response to the medium dose, and 5 in response to the high dose (**Fig. 2G-2I**), further showing prominent species-specific sensitivity to MEHP. In human cells, MEHP selectively upregulated detoxification and antioxidant genes (*GSTM2, GPX3, PRDX1, PRDX3*) and inflammatory mediators (*IL-6R* and *IL-1R1),* while downregulating other antioxidants (*CAT, PRDX2),* the master regulator of lipid metabolism *(PPARG),* and other inflammatory mediators *(IL-24*, *IL17RC*, *IL17RD*, *IL17RE).* These patterns are consistent with findings from Brassea-Pérez et al. (2024), who found that DEHP increases the expression of phase II detoxification genes (e.g., *GSTM1, MGST1, GPx3*) and inflammatory cytokines (IL-6), while suppressing antioxidant genes (*SOD1, CAT, NRF2*), immune genes (*IL-15, IL-16*) and lipid metabolism regulators (*PPARA, PPARD)* in human skeletal muscle cells. In contrast, the same study showed that DEHP upregulated antioxidant (*SOD3*, *GPX2, GPX8)* and damage repair *(NRF2, OGG1, IL-15)* genes and downregulated phase II detoxification GST isoforms in sea lion muscle cells. In our study, seal fibroblasts upregulated antioxidant (*GPX4)* and immune signaling genes (*IL-17)*, whereas dolphin cells downregulated *ILF3,* but did not induce expression of GSTs. This divergence may reflect evolutionary differences, as many cetaceans have lost or pseudogenized GST isoforms, including *GSTM1*, *MGSTM2, GSTA, and GSTT*, which play a crucial role in xenobiotic metabolism (Danneels et al., 2025; Meyer et al., 2019; Tian et al., 2019). Upregulation of *GSTM2* in human but not marine mammal cells suggests that human cells rely on glutathione-mediated detoxification of phthalates.

**Figure 2.**
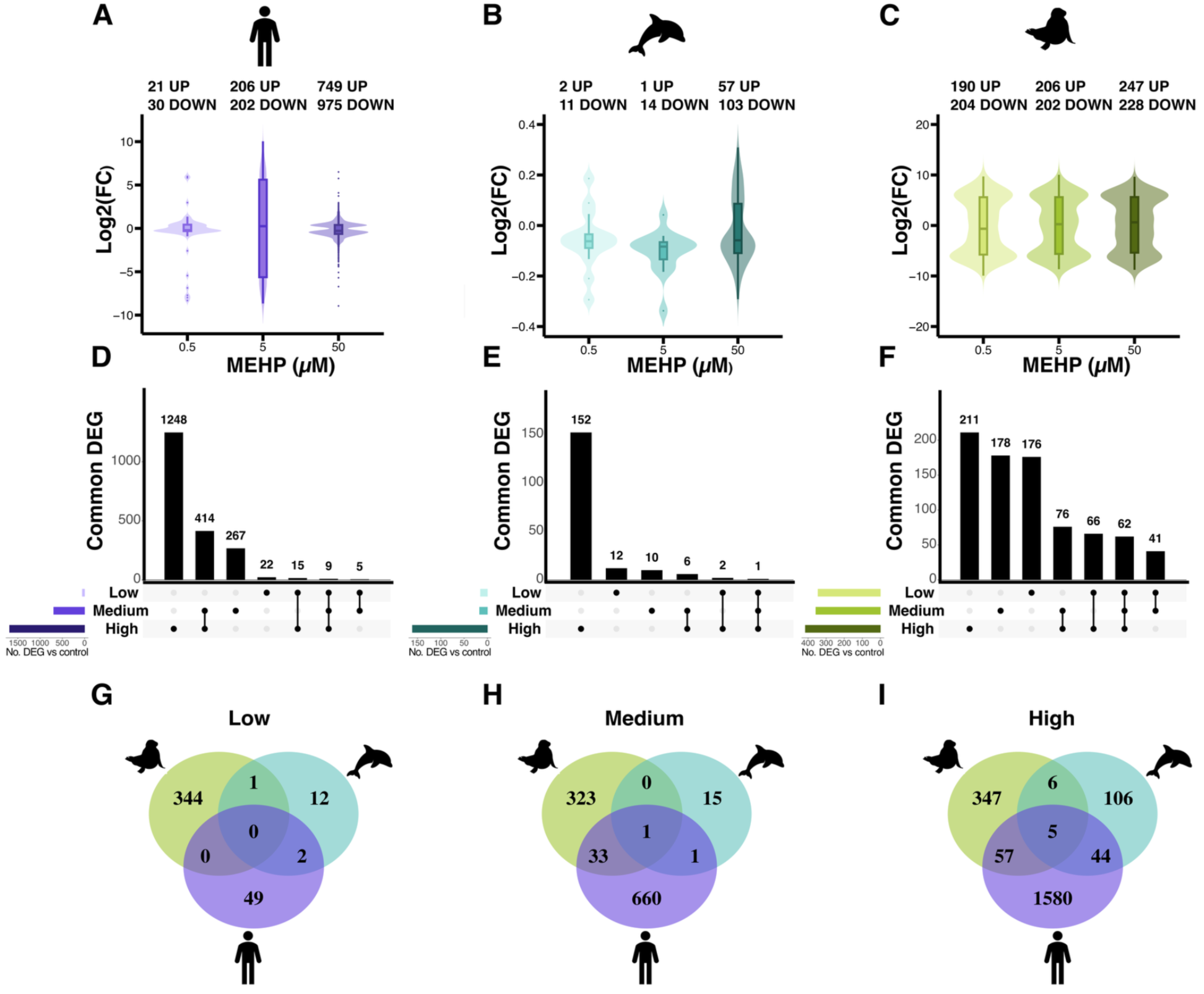
MEHP induces species-specific gene expression signatures in dermal fibroblasts. (A,B,C) Violin plots showing significant (FDR < 0.05) differential gene expression patterns in human (A), dolphin (B), and seal (C) cells following 48 hours of exposure to 0.5 *µ*M, 5 *µ*M, or 50 *µ*M of MEHP. Fold-change values for gene expression were log2-transformed, with positive values indicating upregulation in MEHP-treated cells relative to the control, and negative values representing downregulation. (D,E,F) UpSet plots showing the unique and shared DEGs (FDR < 0.05) found between MEHP treatments for human (D), dolphin (E), and seal cells (F). The bar plots on the left show the total number of DEGs (FDR < 0.05) in either treatment relative to the control. (G,H,I) Venn diagrams showing the number of DEGs (FDR < 0.05) (vs vehicle control) shared between human (G), dolphin (H), and seal cells (I) exposed to low (0.5 *µ*M), medium (5 *µ*M), or high (50 *µ*M) MEHP vs vehicle controls.

Gene Set Enrichment Analysis (GSEA) revealed pathways significantly enriched across all MEHP conditions in each species, except for elephant seal cells at the low dose, despite these cells showing the largest number of DEGs among the three species (**Fig. 3C, 3G**). In the low dose, human cells upregulated genes enriched in energy pathways including glycolysis, oxidative phosphorylation, fatty acid metabolism, and cholesterol homeostasis (**Fig. 3A**). In contrast, dolphin cells upregulated genes enriched in pathways including TNF-α signaling via NFκB, estrogen responses, KRAS signaling, and bile acid metabolism, while downregulating genes enriched in cell cycle regulation and mitotic kinase activity (**Fig. 3D**). At the medium dose, all three species upregulated genes involved in KRAS signaling and TNF-α signaling via NF-κB (**Figs. 3B, 3E, 3G**). Human and seal cells both upregulated genes involved in cell cycle, oxidative phosphorylation, and inflammation (**Figs. 3B, 3G**), while dolphin and human cells shared responses linked to estrogen signaling and hedgehog signaling (**Figs. 3B, 3E**). At this dose, human cells also exhibited the largest number of upregulated pathways (27), including glycolysis, hypoxia, fatty acid metabolism, adipogenesis, tumor suppressor pathways, peroxisome function, cell survival, and immune activation, including IL-2-STAT5 signaling (**Fig. 3B**). In contrast, eight upregulated pathways were enriched in dolphin cells (**Fig. 3E**). In comparison, 12 pathways were enriched in elephant seal cells, which upregulated genes involved in steroid-related processes (*LPIN1, ACE, SIRT1, PIAS2*) and inflammatory signaling (TGF-β, complement) (**Fig. 3G**).

**Figure 3.**
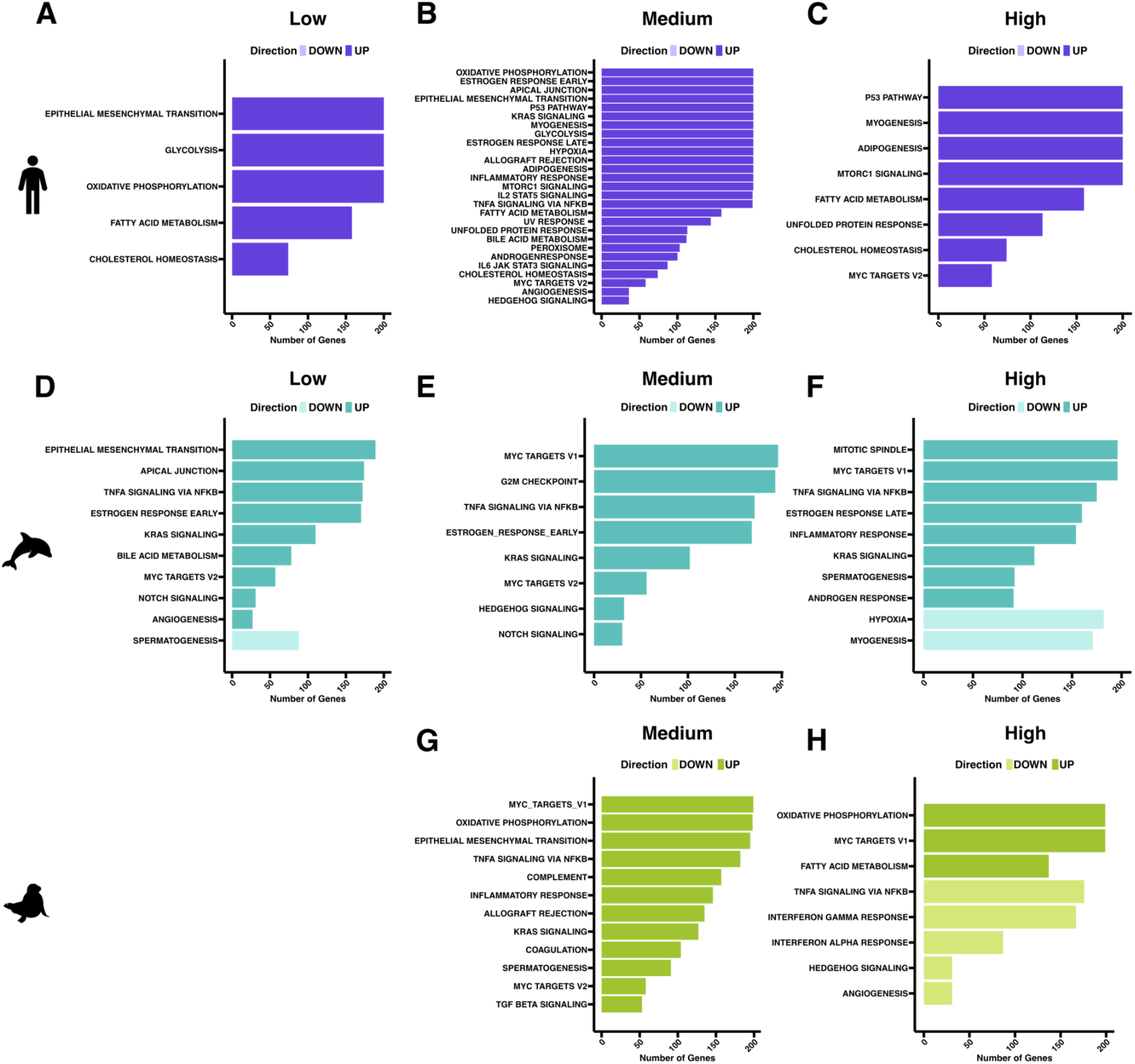
MEHP induces species-specific pathway enrichment. GSEA of biological states or processes (Molecular Signatures Database (MSigDB) significantly (FDR ≤ 0.25) enriched in human (A-C), dolphin (D-F), and seal cells (G, H) exposed to low (0.5 *µ*M), medium (5 *µ*M), or high (50 *µ*M) MEHP levels.

At the high MEHP dose, all three species upregulated genes enriched in cell-cycle pathways (**Figs. 3C, 3F, 3H**). Notably, cells from both marine mammal species exhibited enrichment for hypoxia-related and inflammatory responses, including TNF-α signaling via NF-κB (**Figs. 3F, 3H**), which are crucial for their diving lifestyle (Allen et al., 2024). Moreover, human and seal cells upregulated genes enriched in fatty acid metabolism (**Figs. 3C, 3H**), while dolphin and human cells shared enrichment for genes involved in myogenesis (**Figs. 3C, 3F**). Compared with dolphin and human cells, elephant seal cells upregulated genes enriched for oxidative phosphorylation and downregulated genes enriched for angiogenesis and interferon responses (**Fig. 3H**). In contrast, dolphin cells showed a broader range of enriched pathways, including genes involved in mitosis, estrogen responses, and KRAS signaling (**Fig. 3F**), while human cells showed a unique pattern with genes enriched in cell survival pathways, protein synthesis, unfolded protein response, and lipid metabolism, including cholesterol homeostasis and adipogenesis (**Fig. 3C**). Overall, GSEA revealed dose- and species-specific responses to MEHP. At higher doses, all species upregulated inflammatory and cell-signaling pathways, with human cells showing the broadest response and significant enrichment for energy metabolism, immune activation, and cell-survival pathways. Marine mammal cells exhibited enrichment in hypoxia and inflammation pathways, with species-specific responses: elephant seal cells showed enrichment in metabolism and immune pathways, while dolphin cells upregulated hormone signaling, cell division, and myogenesis.

Bjørneset et al. (2023) reported that killer whale fibroblasts exposed to a mixture of persistent organic pollutants (POPs) downregulated *CYP1A1*, the GR target *ANGPTL4,* immune-related genes (*NR4A3*, *TNFRA5, IL-6*), and *PDK4*, a key regulator of fibroblast metabolism, while upregulating lipid metabolism genes and nuclear receptors (*ADIPOQ*, *CYP4A*, *PPARA*, *PPARG*) at the highest concentration. Similarly, we found that dolphin cells exposed to MEHP downregulated *ANGPTL4*, ACKR3, and CD34, suggesting possible suppression of the glucocorticoid receptor and immune signaling, while elephant seal cells downregulated genes involved in lipid transport and trafficking (EHD1, ABCA2). In human cells, we observed upregulation of genes involved in xenobiotic metabolism and processing (*KCND3*, *CYP19A1*, *CYP1B1*, *SFRP1*, *CYP7B1*) and downregulation of lipid metabolic processes and transport markers (*STC1*, *ANGPTL4*, *ABCA1*, *CD36*, *PDK4*, *FABP4*, *PPARG*). Bjørneset et al. (2023) also found that functional enrichment analysis of DEGs in response to pollutant exposure was strongly associated with striated muscle cell development (*MYH11, OBSCN, MYOM2*) and smooth muscle cell development (*PTAFR, HTR18*). Our analysis revealed significant enrichment of myogenic pathways across species, and upregulation of *MYH11*, a muscle development marker, in elephant seal cells. Differential gene expression in response to MEHP may underlie species-specific variations in energy metabolism, although enrichment for cell cycle, inflammation, and hypoxia-related pathways across all three species, particularly at higher MEHP concentrations, is consistent with the known effects of MEHP (Wang et al., 2012; Manteiga & Lee, 2016; Kim et al., 2023; Kuo et al., 2023). MEHP induces oxidative stress (Brassea-Pérez et al., 2021), mitochondrial dysfunction (Li et al., 2024), and inflammatory signaling through TNF-α/NF-κB (Ashari et al., 2022) in human and rodent models (Tetz et al., 2013; Kocbach Bølling et al., 2011). Upregulation of oxidative phosphorylation and fatty acid metabolism in human and seal cells, but not in dolphin cells, suggests distinct strategies for meeting energy demands and sustaining mitochondrial function under metabolic stress. This is intriguing, as our results show that MEHP suppresses mitochondrial respiration (see Section 3.3) and alters mitochondrial morphology (see Section 3.4) in different ways across the three species.

### 3.3. MEHP Exposure Suppresses Mitochondrial Respiration Across Species

Our RNA-seq results showed enrichment for the two main energy-generating pathways (oxidative phosphorylation and glycolysis) in seal and human cells. Hence, we then assessed mitochondrial function and glycolytic rates using extracellular flux assays. Exposure to higher MEHP concentrations, shown in other studies to disrupt mitochondrial integrity and initiate apoptosis (Erkekoglu and Kocer-Gumusel, 2014), led to a decrease in oxygen consumption rates (OCR) across all species, with more pronounced reductions observed in marine mammal cells compared to human cells (**Figs. 4A, 4D, 4G**). In all species, basal respiration (Human: *P* = 0.002; Dolphin: *P* < 0.0002; Seal: *P* < 0.0005), maximal respiration (Human: *P* = 0.0007; Dolphin: *P* < 0.0001; Seal: *P* < 0.0001), spare respiratory capacity (Human: *P* < 0.0006; Dolphin: *P* < 0.0001); Seal: *P* < 0.0001), and ATP-linked respiration (Human: *P* = 0.001; Dolphin: *P* = 0.0001; Seal: *P* = 0.0002) declined significantly following exposure to the high MEHP dose relative to the control (**Figs. 4B, 4E, 4H**), indicating an overall suppression of mitochondrial metabolism. However, at the low and mid doses, there were minimal changes observed in seal and human cells, with the low and mid dose showing no change relative to the control in human cells (**Figs.4B & 4H**), whereas dolphin cells showed significant reductions in basal respiration (*P* = 0.0009), maximal respiration (*P* = 0.001), spare respiratory capacity (*P* = 0.002), and ATP-linked respiration (*P* = 0.001) at the mid dose relative to the control (**Fig. 4E**). Additionally, elephant seal cells showed significant reductions in proton leak (High: *P* = 0.05) and non-mitochondrial respiration Mid: P = 0.05; High: P = 0.03) relative to the control, but not in dolphin or human cells (Fig. 4H).

Extracellular acidification rate (ECAR) showed no significant change in human cells, even at the highest MEHP dose (**Fig. 4C**). Although mitochondrial respiration was only markedly suppressed at the highest MEHP dose, unchanged ECAR values and increased expression of glycolytic genes (*HK2, G6PD, PFKFB1, ALDOA, ENO1, PGAM1/2, LDHA, LDHC, PKM, PHK1*) in human cells suggest a reliance on glycolytic ATP production and a metabolic shift from oxidative phosphorylation at this dose. This response aligns with the Warburg effect, where glycolytic activity increases in response to mitochondrial dysfunction or cellular stress (Seyfried & Shelton, 2010; Vander Heiden et al., 2009). While glycolysis is less efficient in generating ATP compared to oxidative phosphorylation, this shift reflects an adaptive strategy that enables cells to preserve energy by altering their primary energy source (Epstein et al., 2017).

In dolphin cells, ECAR followed a pattern like that of the OCR (**Fig. 4F**). In contrast, in elephant seal cells, both ECAR and OCR were significantly suppressed at the highest MEHP dose (**Fig. 4I**). Vascular and muscle cells from deep-diving elephant seals regulate mitochondrial dynamics to optimize oxygen use and reduce oxidant generation after extreme hypoxia and in response to other stressors (Allen et al., 2024; Torres-Velarde et al., 2021). In elephant seal cells, mitochondrial respiration appeared to be the preferred energy source until the highest MEHP dose, indicating a delayed or limited shift to glycolysis under MEHP-induced stress. This was accompanied by increased expression of genes involved in oxidative phosphorylation, including key subunits of the electron transport chain (*NDUFA1, NDUFB4, NDUFS1, SDHA, UQCRC1, CYCS, COX4I1, COX5A, ATP5A*), the TCA cycle (*IDH1, IDH3A, OGDH, MDH2, PDHA1, PDK4, MDH1*). Unlike human or seal cells, dolphin cells exhibited significant, dose-dependent reductions in both OCR and ECAR, even at the lowest MEHP concentration, despite showing the most muted response to MEHP in terms of number of DEGs. This was accompanied by upregulation of stress and hypoxia-related genes (*NDUFA4L2, TXNIP, DDIT4, EPAS1*), which suppress mitochondrial respiration and glucose metabolism (Alhawiti et al., 2017; Scortegagna et al., 2003; Michalski et al., 2024; Tello et al., 2011; Torres-Velarde et al., 2021). Similarly, while other studies in cetacean primary fibroblasts confirmed that MEHP acts as a potent nuclear receptor disruptor, they also revealed that cetacean fibroblasts can remain largely unresponsive to direct phthalate exposure (Mukundan et al., 2026, Preprint). Together, these findings show species-specific differences in the regulation of cellular metabolism in response to phthalate exposure, consistent with previous reports indicating that MEHP can impair mitochondrial function by disrupting the electron transport chain, reducing ATP production, and increasing oxidative stress in both human and rodent cells (Rowdhwal and Chen, 2018).

**Figure 4.**
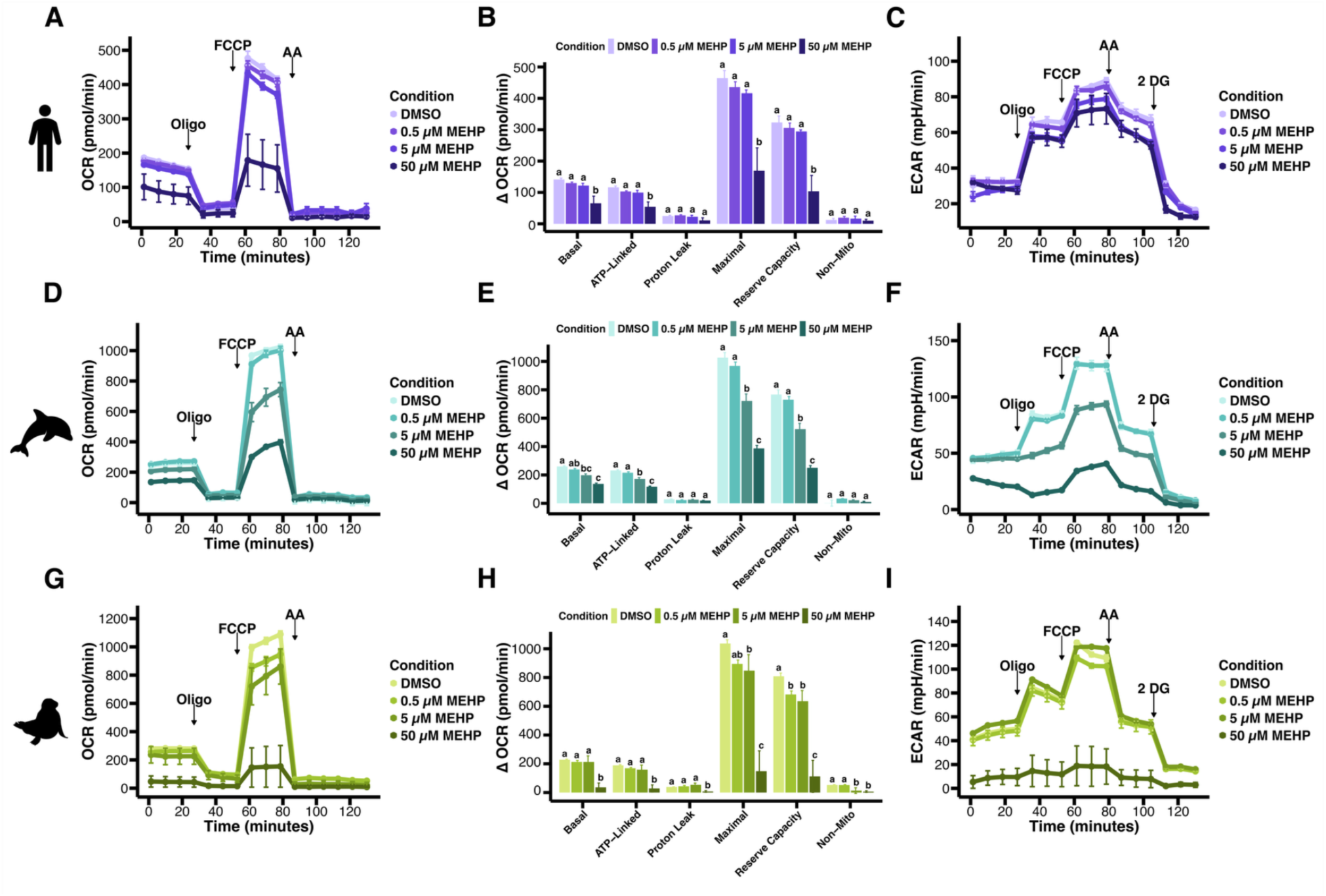
MEHP exposure dysregulates cellular bioenergetics in human, dolphin, and seal cells. (A,D,G) Oxygen consumption rates (OCR), (B,E,H) mitochondrial function, and (C,F,I) extracellular acidification rates in human (A-C), dolphin (D-F), and seal (G-I) fibroblasts. Different letters denote significant differences between conditions. Results are expressed as mean ± SE from n= 3-5 wells per species per condition.

### 3.4. MEHP Exposure Alters Mitochondrial Architecture in Human and Marine Mammal Cells

We next evaluated whether the observed species-specific response to low (0.5 *µ*M) or high (50 *µ*M) MEHP exposure on mitochondrial function (see Section 3.3) were correlated with alterations in mitochondrial morphology using 3D confocal microscopy analysis of the mitochondrial reticulum (**Fig. 5A**). For all three species, MEHP conditions (low and high) did not significantly alter total mitochondrial volume compared to controls (**Fig. 5B**), but did significantly increase mitochondrial disconnections in human (*χ*^2^= 24.54, df = 2, *P* < 0.0001) and seal (*χ*^2^ = 10.32, df = 2, *P* = 0.005) cells at both low (Human: *P* < 0.0001; Seal: *P* = 0.02) and high (Human: *P* = 0.0004; Seal: *P* = 0.02) MEHP doses (**Fig. 5B**). Additionally, Mnf2, which is essential for mitochondrial fusion (Santel et al., 2003), significantly decreased in human (F_2,6_ = 34.94, *P* = 0.0005) cells at low (*P* = 0.002) and high (*P* = 0.0005) MEHP doses, but not in seal or dolphin cells (**Fig. 5C**). In humans, this increase in mitochondrial fragmentation and decrease in mitochondrial fusion protein Mfn2, was associated with suppressed respiration (see Section 3.3). Loss of Mfn2, a protein that induces fusion of the outer mitochondrial membrane, disrupts mitochondrial membrane potential and promotes mitochondrial fragmentation in mammalian cells (Santel et al., 2003). Furthermore, this response may reflect a coordinated but ultimately overwhelmed metabolic defense mechanism leading to structural collapse of the mitochondrial network in human cells exposed to phthalates. Although mitochondrial fragmentation increased in seal cells, Mfn2 protein levels did not significantly change; however, we observed a distinct upward trend in response to phthalate exposure. This trend was mirrored in our transcriptomic analysis, with significant increases in the expression of genes encoding mitochondrial fusion/fission and trafficking proteins (*OPA1, MFN2, RHOT1, RHOT2*). This response might reflect an evolutionary adaptation to maintain mitochondrial function in energy-challenging conditions such as breath-hold diving (Kooyman and Ponganis, 1998; Davis, 2013). In contrast, the absence of mitochondrial fragmentation and fusion changes despite functional decline in dolphin cells further suggests distinct, species-specific cellular structural and functional responses to phthalate exposure. These findings suggest higher metabolic plasticity in dolphin cells under phthalate-induced stress, potentially reflecting the fully aquatic phenotype (Hindle, 2020). Overall, these results show species-specific changes in mitochondrial architecture in response to phthalate exposure. Furthermore, these results suggest that the observed dysregulation in mitochondrial function may be related to fragmentation of mitochondrial reticulum in human and seal cells.

**Figure 5.**
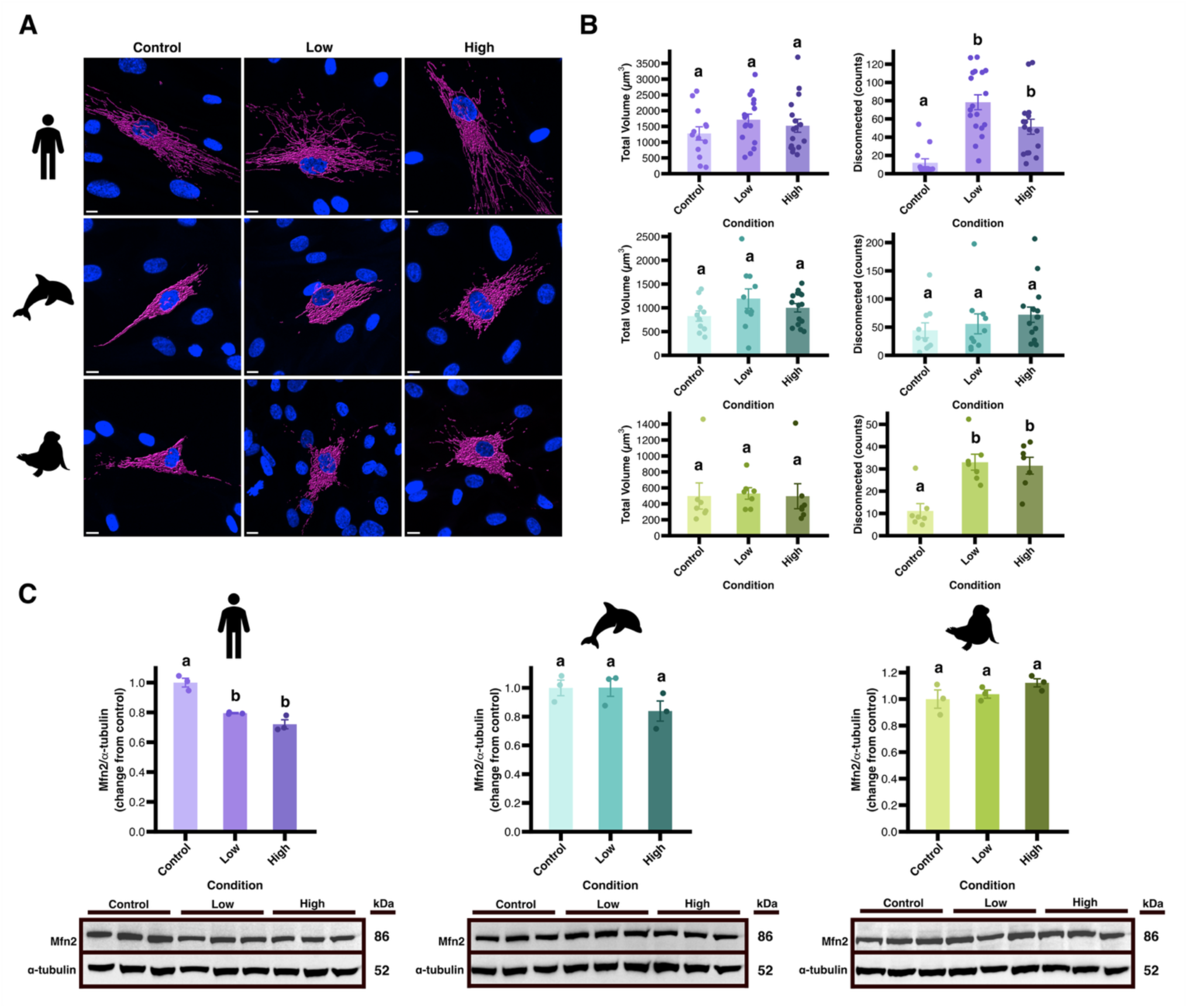
MEHP exposure alters species-specific mitochondrial architecture. (A) 3D confocal reconstruction visualizing mitochondrial morphology in seals exposed to phthalates (scale bar is 10 *µ*m), (B) analysis of mitochondrial networks, and (C) WB analysis of Mfn2 in human, dolphin, and seal fibroblasts following 48 hour exposure to low (0.5 *µ*M) and high (50 *µ*M) MEHP. Different letters denote significant differences between conditions. Results are expressed as mean ± SE.

Overall, our study shows that MEHP induces species-specific transcriptional and bioenergetic signatures in cells from marine mammals and humans. While MEHP did not induce cytotoxicity, it upregulated cell cycle pathways, altered mitochondrial respiration and architecture, and activated metabolic and inflammatory signaling pathways. Human cells showed the most pronounced changes in gene expression, while dolphin cells exhibited the greatest bioenergetic plasticity. These findings suggest that species-specific metabolic adaptations may influence cellular responses to MEHP. The variability in responses underscores the need for species-specific considerations in environmental risk assessments. While marine mammals may benefit from protective adaptations, chronic exposure to MEHP could still compromise key processes, such as mitochondrial function and immune regulation, with potential long-term health impacts. Future research should investigate the effects of MEHP on other cell types and explore mitochondrial dynamics to deepen our understanding of its impact on cellular integrity across species. In this study, the use of cellular models enabled investigation of species-specific global gene expression and bioenergetic responses to chemical pollutants in species for which experimental manipulation is infeasible (Vazquez et al., 2024)

## AUTHOR INFORMATION

### Author Contributions

E.R.P., A.G., and J.P.V-M conceived and designed the research. E.R.P. and J.P.V-M. drafted the manuscript. E.R.P. prepared figures and conducted all experiments. E.K.L, K.N.A., J.P.V-M. and D.E.C. conducted field work. E.R.P, E.K.L and K.N.A. generated cell lines. D.D.M-S conducted the bioinformatic analysis. E.R.P., E.K.L., D.D.M-S., K.N.A., D.E.C., A.G. and J.P.V-M edited and revised the manuscript. All authors have approved the final version of the manuscript and agree to be accountable for all aspects of the work, ensuring that questions regarding the accuracy or integrity of any part of the work are appropriately investigated and resolved. All people designated as authors qualify for authorship, and all who qualify are listed.

## ACKNOWLEDGMENTS

ERP is supported by the Office of Naval Research (ONR) under award number NDSEG10994BIOSCI. JPV-M is supported by R35GM146951. Research funded by the UC Berkeley Peder Sather Center and the Research Council of Norway grant #334739. We thank Dr. Dianna Xing for her assistance in setting up the bioenergetic experiments; Barbie Halaska and the rest of the TMMC staff for the dolphin tissue samples. Special thanks to the RCNR Biological Imaging Facility at UC Berkeley, specifically Facility Director Dr. Denise Schichnes and Facility Scientist Dr. Juliana Cho, for their essential training and support in confocal imaging and analysis. The Vázquez-Medina Lab recognizes that Berkeley sits on the territory of Xučyun, the ancestral and unceded land of the Chochenyo Ohlone, the successors of the historic and sovereign Verona Band of Alameda County. This land was and continues to be of great importance to the Ohlone people. We recognize that every member of the Berkeley community has, and continues to benefit from the use and occupation of this land, since the institution’s founding in 1868. Consistent with our values of community and diversity, we have a responsibility to acknowledge and make visible the university’s relationship to Native peoples. By offering this Land Acknowledgement, we affirm Indigenous sovereignty and will work to hold University of California, Berkeley, more accountable to the needs of American Indian and Indigenous peoples.

